# AAV-Mediated Astrogliosis Recapitulates Early Neuroinflammatory and Bioenergetic Features of Alzheimer’s Disease in Mice

**DOI:** 10.64898/2026.11.18.767679

**Authors:** Akila Ramesh, Narayan Datt Soni, Dipak Roy, Anshuman Swain, Sunil Kumar Khokhar, Paul Jacobs, Abeer Mathur, Halvor Juul, Dushyant Kumar, Ravi Prakash Reddy Nanga, David A Wolk, Virginia MY Lee, John A Detre, Ravinder Reddy, Mohammad Haris

**Author notes:** Address for Correspondence, Mohammad Haris, Ph.D., Center for Advanced Metabolic Imaging in Precision Medicine, 422 Curie Boulevard, B1, Stellar Chance Labs, Philadelphia, PA 19104-6100. Akila Ramesh and Narayan Datt Soni contributed equally to this work.

## Abstract

Neuroinflammation and altered neuron-astrocyte metabolic interactions are increasingly recognized as early features of Alzheimer’s disease (AD). However, the underlying metabolic adaptations remain incompletely understood. This study employed deuterium magnetic resonance spectroscopy (DMRS) following administration of deuterium labeled glucose and acetate to investigate neuronal and astroglial substrate utilization and metabolic fluxes, respectively, in adenovirus-induced neuroinflammation and early-stage APP^NL-F/NL-F^ mice. In adenovirus-induced astrogliosis, increased acetate derived [4,4’-^2^H_2_]Glx (d_2_-Glx) production suggested enhanced astrocytic acetate utilization and TCA cycle activity. Conversely, reduced glucose derived d_2_-Glx despite comparable glucose uptake suggested a diminished neuronal activity due to potentially impaired neuronal oxidative metabolism. However, the glucose-derived lactate labeling remained unchanged, suggesting differential effects on oxidative and non-oxidative pathways of glucose metabolism. Similarly, 3-month-old APP^NL-F/NL-F^ mice exhibited significantly increased [2,2’,2’’-^2^H_3_]acetate (d_3_-Ace) influx compared with wild-type controls, suggesting early alterations in astrocytic substrate utilization. While glucose uptake remained unchanged, glucose derived Glx and lactate showed modest, non-significant decreases, potentially reflecting early alterations in glucose metabolism. Overall, both models exhibited a metabolic pattern characterized by increased astrocytic acetate utilization with trends towards reduced glucose associated metabolism, with more pronounced changes in adenovirus-induced astrogliosis. These findings highlight the utility of DMRS for investigating dynamic metabolic adaptations during neuroinflammation and early AD pathogenesis. Importantly, the adenovirus-induced astrogliosis model recapitulates key metabolic alterations associated with the neuroinflammatory aspects of AD. As such, it provides a valuable complementary platform for investigating mechanisms driving early metabolic dysfunction and evaluating therapeutic strategies aimed at restoring neuron-astrocyte metabolic interactions prior to the onset of substantial neurodegeneration.

## 1. Introduction

Alzheimer’s disease (AD) is a progressive neurodegenerative disorder characterized by amyloid-β deposition, tau pathology, synaptic dysfunction, and cerebral glucose hypometabolism (Bano et al., 2023; Scheltens et al., 2016). Emerging evidence suggests that these pathological alterations interact with and aggravate neuroinflammatory pathways, forming a self-reinforcing cycle that drives neuronal dysfunction, cognitive decline, and dementia (Leng & Edison, 2021). Although substantial progress has been made in understanding amyloid-associated pathology, alterations in cellular metabolism and interactions between astrocytes and neurons are increasingly recognized as important aspects of disease progression (Hark & Savas, 2021; Patel et al., 2018). Neuroinflammation is particularly relevant, as genetic and experimental evidence implicates innate immune mechanisms and glial dysfunction in the pathogenesis of sporadic AD (Heneka et al., 2015).

Glutamate (Glu) serves as a major excitatory neurotransmitter in the mature mammalian brain. It is primarily synthesized *de novo* from a tricarboxylic acid cycle (TCA) intermediate, α-keto-glutarate (α-KG), and *via* glutamate-glutamine (Glu-Gln) cycling between neurons and astrocytes. Astrocytes play essential roles in maintaining cerebral energy homeostasis, supporting neuronal function, and regulating neurotransmitter metabolism through the glutamate-glutamine cycle (Andersen & Schousboe, 2023). Alterations in astroglial and neuronal metabolic activity have been demonstrated in experimental AD models, with increased astroglial and reduced neuronal metabolism alongside impaired neurotransmitter cycling (Patel et al., 2018).

Given the pivotal role of neuroinflammation in AD, it is important to study its metabolic consequences in the brain, which can be used to devise a therapeutic strategy aimed at reversing these alterations. Interestingly, such studies have been conducted primarily in amyloid- and tau-based transgenic animal models of AD, in which neuroinflammatory pathways are activated as a consequence of amyloid and tau pathology. On the other hand, the majority of AD cases are sporadic, arising from a complex interplay of aging, genetic susceptibility, and environmental and metabolic risk factors rather than mutations in familial AD-associated genes (Lane et al., 2018). Consequently, investigating the pathological mechanisms underlying sporadic AD is essential for understanding the broader clinical spectrum of the disease. Experimental models targeting individual pathological processes, such as neuroinflammation, therefore provide useful approaches for investigating mechanisms that may contribute to sporadic AD.

Deuterium magnetic resonance spectroscopy (^2^H-MRS/DMRS) enables detection of deuterium-labeled substrates and their downstream metabolic products, allowing monitoring of cell-specific metabolic activity following administration of labeled glucose and acetate (De Feyter et al., 2018; De Feyter & de Graaf, 2021). Acetate is preferentially transported and metabolized by astrocytes, making labeled acetate a useful tracer for investigating astroglial metabolism via label accumulation in glutamine (Waniewski & Martin, 1998). In contrast, glucose-derived labeling reflects overall cerebral glucose metabolism and primarily reflects neuronal metabolic activity via glutamate labeling, given that neurons account for the majority of oxidative glucose consumption (De Feyter & de Graaf, 2021; Patel et al., 2018). Although ^2^H-MRS cannot resolve glutamate and glutamine resonances *in vivo,* which overlap as a composite Glx peak, ^2^H-labeled Glx derived from labeled glucose and acetate can serve as practical proxies for neuronal and astroglial metabolism, respectively. Despite this limited spectral resolution, lower gyromagnetic ratio (6.54 MHz/T), and intrinsic MR sensitivity of ^2^H, the practical advantages of ^2^H-MRS are substantial. Specifically, the short longitudinal relaxation time (T_1_) of ^2^H enables rapid signal averaging, achieving adequate SNR within clinically feasible scan times. Moreover, its low natural abundance (0.01156%) renders endogenous metabolite background signals virtually undetectable, while naturally abundant semi-heavy water (HDO; ∼10.12 mM in brain tissue) provides a convenient internal concentration reference for absolute quantification (Soni et al., 2023). Furthermore, recent clinical studies have demonstrated the feasibility of deuterium metabolic imaging (DMI) for detecting altered glucose-derived metabolic fluxes in patients with Alzheimer’s disease (AD) (Khan et al., 2024).

The APP^(NL-F/NL-F)^ mouse model carries humanized familial AD-associated mutations in the amyloid precursor protein gene without APP overexpression (Saito et al., 2014). Notably, hippocampal and cortical proteomic alterations have been identified in these mice as early as three months of age, preceding amyloid plaque development and overt memory impairment (Schedin-Weiss et al., 2020). These findings suggest that molecular and functional disturbances may emerge during early pathological stages, before the onset of established amyloid deposition.

In the present study, we utilized an adenovirus-induced neuroinflammation mouse model to measure ^2^H-labelled glucose and acetate metabolism as proxies for neuronal and astroglial activity, respectively, using ^2^H-MRS. Subsequently, these metabolic measures were investigated in three-month-old APP^(NL-F/NL-F)^ mice to characterize early amyloid-associated metabolic alterations. By examining these complementary models, this study aims to elucidate the metabolic consequences of neuroinflammation and early amyloid-associated pathology and provide insights into astroglial-neuronal metabolic dysfunction relevant to AD.

## 2. Materials and Methods

### 2.1. Animal Models and Experimental Design

All experimental procedures were reviewed and approved by the Institutional Animal Care and Use Committee (IACUC) at the University of Pennsylvania. Six-month-old male C57BL/6 mice (n = 18) were randomly assigned to one of two experimental groups: (i) astrogliosis mice treated with vehicle (AG+VEH, n = 9), (ii) control mice treated with vehicle (CON+VEH, n = 9). Additionally, 3-month-old male APP ^NL-F/NL-F^ (n = 6) and age-matched wild-type (WT; n = 6) mice were used in this study.

### 2.2. Stereotaxic Viral Injections

Mice were anesthetized using isoflurane in Air: O_2_ (70:30) and positioned in a stereotaxic apparatus (Stoelting Co., USA). Bupivacaine (2 mg/kg) was administered subcutaneously over the skull as a local anesthetic prior to the surgical procedure. Following a midline scalp incision, the skull was exposed, and the hippocampal CA1 region was located using bregma as the anatomical reference point. The following stereotaxic coordinates were used for bilateral injections: anteroposterior (AP), −2.0 mm; mediolateral (ML), ±2.0 mm; and dorsoventral (DV), −1.4 mm relative to bregma (Nam et al., 2023). Small burr holes were drilled bilaterally on the skull at the designated coordinates for needle insertion. Ad5-GFAP-EGFP adenoviral particles (1.03 × 10^12^ VP/mL; Creative Biogene) were administered using a 32-gauge Hamilton syringe. A total volume of 1 µL was injected into the hippocampal CA1 region at a constant infusion rate of 0.15 µL/min (∼7 min). Following completion of the injection, the needle was maintained in position for an additional 7 min to facilitate local diffusion of the viral suspension and minimize backflow along the injection tract. The same procedure was subsequently repeated in the contralateral side. Following viral administration, the scalp incision was sutured, and Betadine (povidone-iodine) was applied. Throughout the surgical procedure and recovery period, body temperature of mice was maintained at 37°C using a temperature-controlled heating system. Animals were monitored until recovery from anesthesia and then housed individually until the scheduled DMRS scanning session.

### 2.3. In Vivo Deuterium Magnetic Resonance Spectroscopy (DMRS)

Deuterium magnetic resonance spectroscopy (^2^H-MRS) was performed on a 30 cm diameter horizontal-bore 9.4 T scanner equipped with Avance III HD console and ParaVision 6.0.1 software (Bruker BioSpin, Germany). Mice were fasted for 6-12 h prior to scanning and anesthetized with 1.5–2% isoflurane in Air: O_2_ (70:30). The head was placed inside a custom-built head cone pre-mounted with a home built four turns ^2^H transmit/receive surface coil. This assembly was placed inside a 20 mm dual channel ¹H transceiver volume coil (m2m Imaging, USA) with ^2^H surface coil and ^1^H volume coil centers aligned. Finally, the volume coil was placed inside the magnet and centers of coils were matched with magnet isocenter. Throughout the experiment, core body temperature was maintained at 37°C, and respiration was continuously monitored and regulated to maintain a rate of 60-80 breaths per minute.

A baseline ^2^H-MRS spectrum was acquired prior to substrate administration using a pulse-acquire sequence: 750 Hz bandwidth, 65° flip angle, 128 points, 2,048 averages, and repetition time (TR) of 200 ms. To assess substrate-specific cerebral metabolic kinetics, mice received a 90-minute subcutaneous infusion of either [2,2’,2’’-^2^H_3_]acetate (d_3_-acetate: 1 mol/L in H_2_O, pH 7) or [6,6’-^2^H_2_]glucose (d_2_-glucose: 1 mol/L in H_2_O, pH 7), administered using a modified bolus-variable-rate infusion protocol (Fitzpatrick et al., 1990; Soni et al., 2023). Infusion rates were optimized to achieve steady-state substrate concentrations in brain tissue. Following initiation of the infusion, 13 subsequent ^2^H-MR spectra were continuously acquired using same parameters at ∼7-minute intervals, yielding a time-resolved dataset of cerebral substrate uptake and metabolic turnover over the 90-minute infusion period.

### 2.4. Spectral Analysis

Raw free induction decays (FIDs) were processed using Mnova 14.3.0 (Mestrelab Research, Santiago de Compostela, Spain). Preprocessing consisted of phase correction, baseline correction, and zero-filling to extend the data points from 256 to 512. FIDs were transformed into the frequency domain using a 5-Hz exponential line-broadening filter. To optimize the signal-to-noise ratio (SNR), downfield ^2^H spectra underwent singular value decomposition (SVD) denoising applied to a Casorati matrix (m x n: 14 x 513). The dominant singular values corresponding to the expected post-infusion metabolic resonance signals were selected for matrix reconstruction, yielding an approximate five-fold increase in baseline SNR.

Resonance signals were fitted with Lorentzian line shapes using the Lmfit package in Python (version 3.13.14). Fitted peak integrals for glucose (Glc), acetate (Ace), lactate (Lac), and glutamate/glutamine (Glx) were resolved and their concentrations were derived relative to the baseline HDO signal. The concentration of naturally abundant HDO was defined as 10.12 mM based on the established tissue water fraction in the mouse brain. To correct for T_1_ saturation effects, T_1_ relaxation times for each metabolite were determined from respective phantoms at 37°C on a 9.4-T Bruker Avance III magnet using an inversion recovery sequence (12 inversion times logarithmically spaced from 10 to 3000 ms; repetition time (TR) = 3000 ms).

To calculate the initial flux rates from the metabolite turnover curves, we applied the following equation:

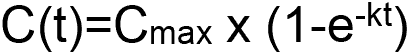

Where C(t) represents the labeled metabolite concentration in mM at any given time point t (min), C_max_ is the estimated maximum steady-state plateau concentration in mM, and k is the rate constant (min^-1^).

### 2.5. Statistical Analysis

Statistical analyses were performed using GraphPad Prism software (Version 8.0; GraphPad Software, San Diego, CA, USA). The normality of data distribution was evaluated using the Shapiro-Wilk test. For non-normally distributed data, a two-tailed Mann-Whitney U test was employed to assess the statistical significance of differences in steady-state concentrations and initial rates between the experimental groups. A p-value less than 0.05 was considered statistically significant. All data are presented as mean ± standard deviation (SD).

## 3. Results

### 3.1. Astroglial and Neuronal Metabolism in Adenovirus Treated Mice

#### 3.1.1. Acetate Metabolism

Following d_3_-acetate infusion, its level in the brain increased with time, achieving a steady-state plateau concentration at around 60 min **(Fig. 1A)**. The absolute steady-state concentration of d_3_-acetate did not vary significantly between the groups (p=0.69), although astrogliosis mice exhibited a slightly higher d_3_-acetate labeling (AG: 2.27±0.40 mM) compared to controls (CON: 2.08±0.23 mM) **(Fig. 2C)**. Fitting the measured values to an exponentially saturating curve demonstrated that the initial rate of acetate labeling was relatively higher in the AG group (0.106±0.036 μmol/g/min) relative to the CON group (0.084±0.024 μmol/g/min), though this difference did not cross the threshold of statistical significance (p=0.49, Fig. 2A).

**Figure 1.**
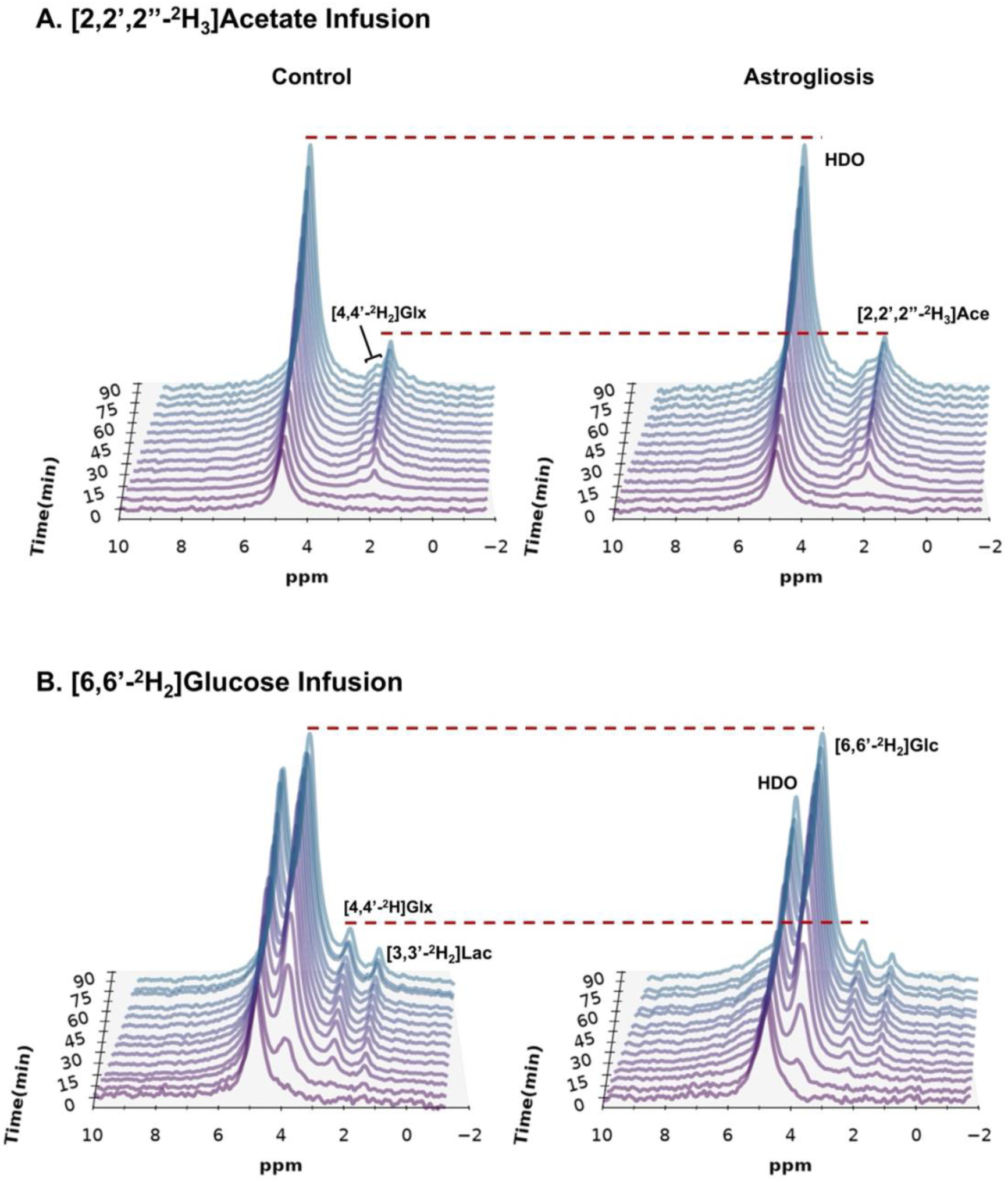
Time series plot of denoised ^2^H spectra from astrogliosis (right) and control (left) mice obtained with a temporal resolution of 6.83 minutes, **(A)** shows the evolution of HDO (∼4.8 ppm), [2,2’,2’’-^2^H_3_]Ace (∼2.0 ppm), and [4,4’-^2^H_2_]Glx (∼2.4 ppm) following subcutaneous [2,2’,2’’-^2^H_3_]acetate administration. **(B)** depicts HDO (∼4.8 ppm), [6,6’-^2^H_2_]glucose (∼ 3.8 ppm), [4,4’-^2^H]Glx (∼2.4 ppm) and [3,3’-^2^H_2_]Lactate (∼1.4 ppm) peaks following subcutaneous [6,6’-^2^H_2_]glucose administration. Abbreviations used are: [2,2’,2’’-^2^H_3_]Ace, [2,2’,2’’-^2^H_3_]acetate; [6,6’-^2^H_2_]Glc, [6,6’-^2^H_2_]glucose; [4,4’-^2^H]Glx, [4,4’-^2^H]glutamate/glutamine; [3,3’-^2^H_2_]Lac, [3,3’-^2^H_2_]lactate.

**Figure 2.**
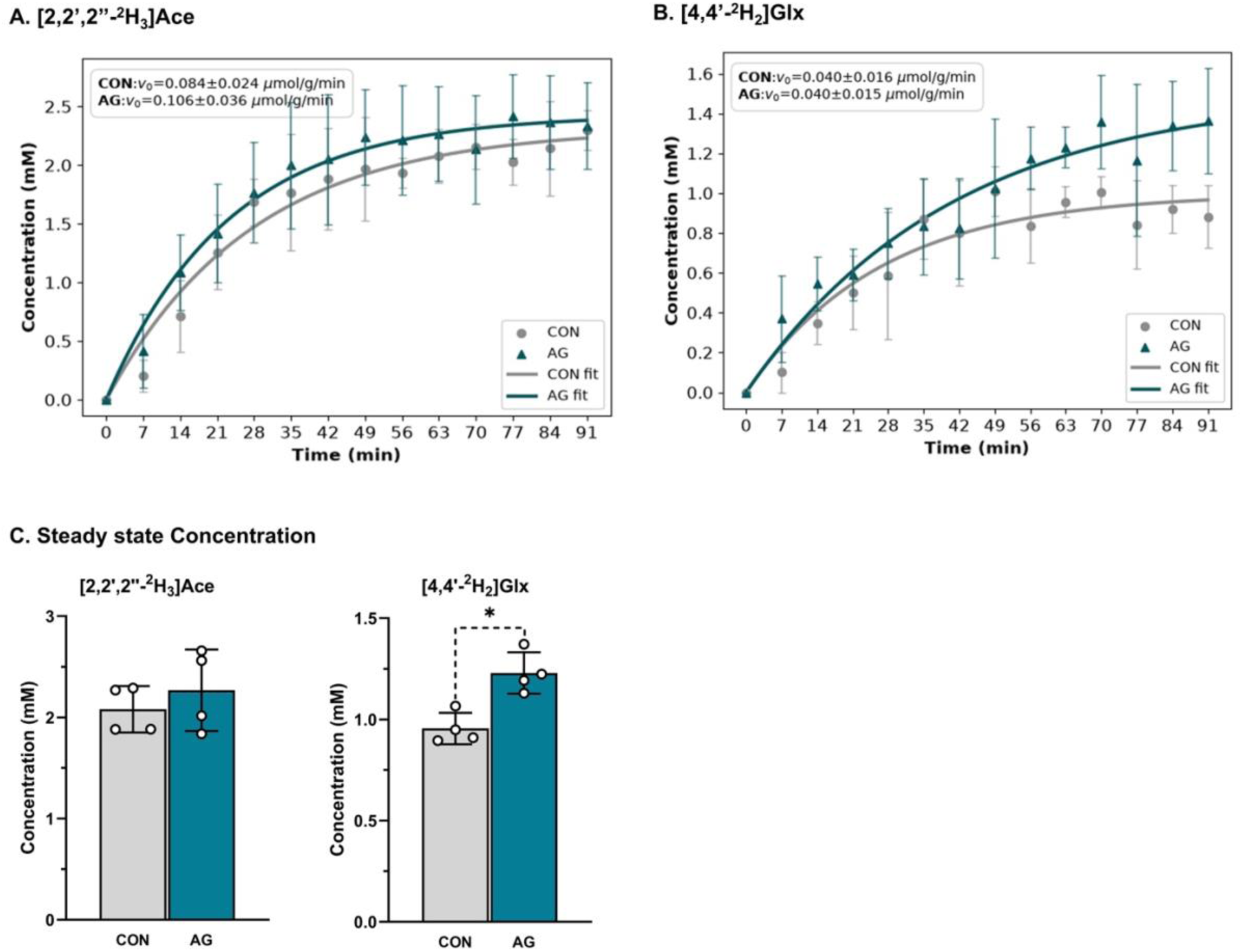
Turnover curves showing concentrations of A. [2,2’,2’’-^2^H_3_]Ace, B. [4,4’-^2^H_2_]Glx in the brains of CON (grey circles) and AG (blue triangles) as a function of time. The symbols represent mean values at each timepoint, and solid lines represent exponential fits to the data. C. Steady state (∼60min) concentrations of [2,2’,2’’-^2^H_3_]acetate and [4,4’-^2^H_2_]Glx represented as means±SD, *p<0.05. Abbreviations used are: [2,2’,2’’-^2^H_3_]Ace, [2,2’,2’’-^2^H_3_]acetate; [6,6’-^2^H_2_]Glc, [6,6’-^2^H_2_]glucose; [4,4-^2^H_x_]Glx, [4,4-^2^H_x_]glutamate/glutamine; [3,3-^2^H_2_]Lac, [3,3-^2^H_2_]lactate, AG, astrogliosis; CON, control.

Downstream metabolism through the astrocytic tricarboxylic acid (TCA) cycle was monitored via the synthesis of [4,4’-^2^H_2_]-glutamate/glutamine (d_2_-Glx). The steady-state concentration of d_2_-Glx derived from acetate achieved a significantly higher (1.23±0.10 mM) level when compared to CON mice (0.95±0.08 mM; p=0.03) **(Fig. 2C)**. In contrast, the initial rates of d_2_-Glx labeling were indifferent across two groups (CON: 0.038±0.005 μmol/g/min vs. AG: 0.037±0.003 μmol/g/min; p>0.99) **(Fig. 2B)**.

#### 3.1.2. Glucose Metabolism

To determine whether adenovirus-mediated neuroinflammation compromised primary neuronal bioenergetics or triggered counter-regulatory shifts in glucose handling, DMRS metrics were tracked concurrently during the infusion of d_2_-glucose. The d_2_-glucose reached a stable steady state concentration at roughly 60 min post-infusion (Fig. 3A). The steady-state concentration of d_2_-glucose were comparable between the groups (AG: 7.21±0.73 mM vs. CON: 7.17±0.71 mM; p=0.84) **(Fig. 3D)**. Similarly, the initial rates of glucose uptake in AG mice (0.267±0.032 μmol/g/min) closely matched the rates of CON mice (0.277±0.073 μmol/g/min; p=0.84) **(Fig. 3A)**. The AG mice exhibited a significant reduction in steady-state glucose-derived d_2_-Glx labeling (1.17±0.33 mM) when compared with CON mice (1.78±0.32 mM, p=0.03) **(Fig. 3D)**. Accordingly, the initial flux rate of d_2_-Glx labeling showed a downward trend in the AG group (0.042±0.005 μmol/g/min) than in the CON (0.066±0.005 μmol/g/min, p=0.06) **(Fig. 3B)**. The steady-state concentrations of [3,3’-^2^H_2_]-lactate (d_2_-lactate) in AG mice was found to be 0.32±0.07 mM, which was comparable (p=0.31) to the baseline of 0.37±0.06 mM in the CON **(Fig. 3D)**. The calculated initial rates for the formation of d_2_-lactate were indifferent between the groups (AG: 0.016±0.007 μmol/g/min vs. CON: 0.016±0.007 μmol/g/min; p>0.99) **(Fig. 3C)**.

**Figure 3.**
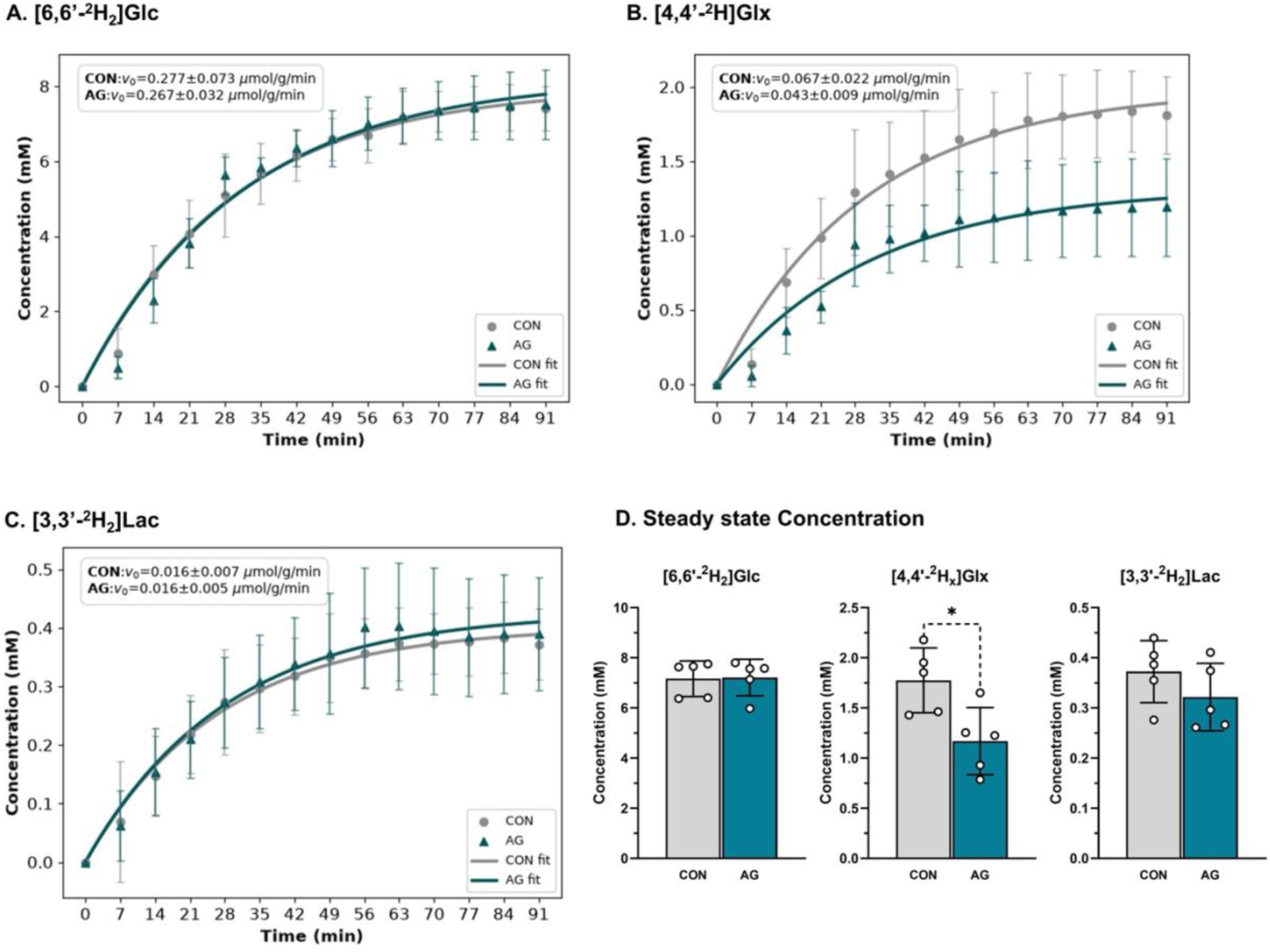
Turnover curves showing concentrations of A. [6,6’-^2^H_2_]Glc, B. [4,4’-^2^H]Glx and C.[3,3’-^2^H_2_]Lac in the brains of CON (grey circles) and AG (blue triangles) as a function of time. The symbols represent mean values at each timepoint, and solid lines represent exponential fits to the data. D. Steady state (∼60min) concentrations of [6,6’-^2^H_2_]glucose, [4,4’-^2^H]Glx, and [3,3’-^2^H_2_]Lac represented as means±SD, *p<0.05. Abbreviations used are: [2,2’,2’’-^2^H_3_]Ace, [2,2’,2’’-^2^H_3_]acetate; [6,6’-^2^H_2_]Glc, [6,6’-^2^H_2_]glucose; [4,4’-^2^H]Glx, [4,4’-^2^H]glutamate/glutamine; [3,3’-^2^H_2_]Lac, [3,3’-^2^H_2_]lactate, AG, astrogliosis; CON, control.

### 3.2. Astroglial and Neuronal metabolism in in APP ^NL-F/NL-F^ Mice

#### 3.2.1. Acetate Metabolism

The metabolic turnover curves were evaluated in 3-month-old APP ^NL-F/NL-F^ (AD) mice and age-matched wild-type (WT) controls. The initial rate of d_3_-acetate influx was significantly elevated in the AD cohort (0.098±0.012 μmol/g/min) compared to WT counterparts (0.049±0.005 μmol/g/min; p=0.05) **(Fig. 4A)**. Similarly, the steady-state acetate pool within AD brains (1.54±0.40 mM) was slightly elevated relative to WT controls (1.33±0.21 mM), though the shift did not achieve statistical significance (p=0.7; Fig. 4C).

**Figure 4.**
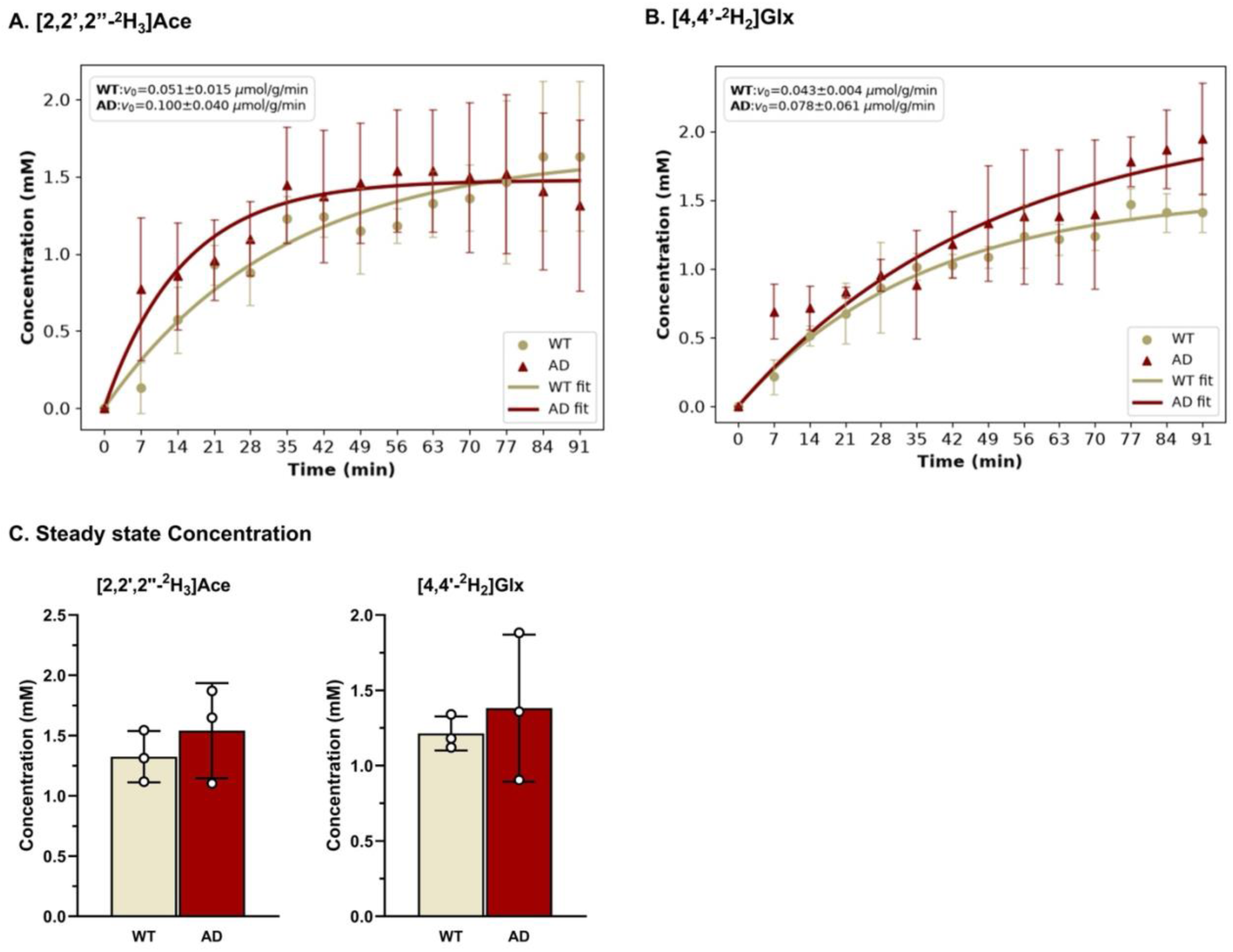
Turnover curves showing concentrations of A. [2,2’,2’’-^2^H_3_]Ace, B. [4,4’-^2^H_2_]Glx in the brains of wild-type (WT) (ivory circles) and APP ^NL-F/NL-F^ Alzheimer’s disease (AD) mice (red triangles) as a function of time. The symbols represent mean values at each timepoint, and solid lines represent exponential fits to the data. C. Steady state (∼60min) concentrations of [2,2’,2’’-^2^H_3_]acetate and [4,4’-^2^H_2_]Glx represented as means±SD. Abbreviations used are: [2,2’,2’’-^2^H_3_]Ace, [2,2’,2’’-^2^H_3_]acetate; [6,6’-^2^H_2_]Glc, [6,6’-^2^H_2_]glucose; [4,4’-^2^H_2_]Glx, [4,4’-^2^H_2_]glutamate/glutamine; [3,3’-^2^H_2_]Lac, [3,3’-^2^H_2_]lactate.

The initial rates of d_2_-Glx labeling from d_3_-acetate were not different between both groups (WT: 0.042±0.002 μmol/g/min vs. AD: 0.043±0.006 μmol/g/min; p=0.35). At the 60-minute mark, the final steady-state concentration of d_2_-Glx in the AD group (1.38±0.49 mM) was very similar to WT mice (1.21±0.11 mM, p=0.70) **(Fig. 4C)**.

#### 3.2.2. Glucose Metabolism

In contrast to the observed changes in acetate metabolism, baseline glucose delivery and accumulation remained preserved in the AD mice. The steady-state concentration of cerebral d_2_-glucose in AD mice (6.45±0.74 mM) was statistically comparable to that of WT control mice (6.01±0.69 mM; p=0.40) **(Fig. 5D)**. The initial rates of glucose uptake remained unchanged between two groups, with AD animals exhibiting a rate of 0.215±0.014 μmol/g/min compared to 0.219±0.014 μmol/g/min in WT controls (p=0.20) **(Fig. 5A)**.

**Figure 5.**
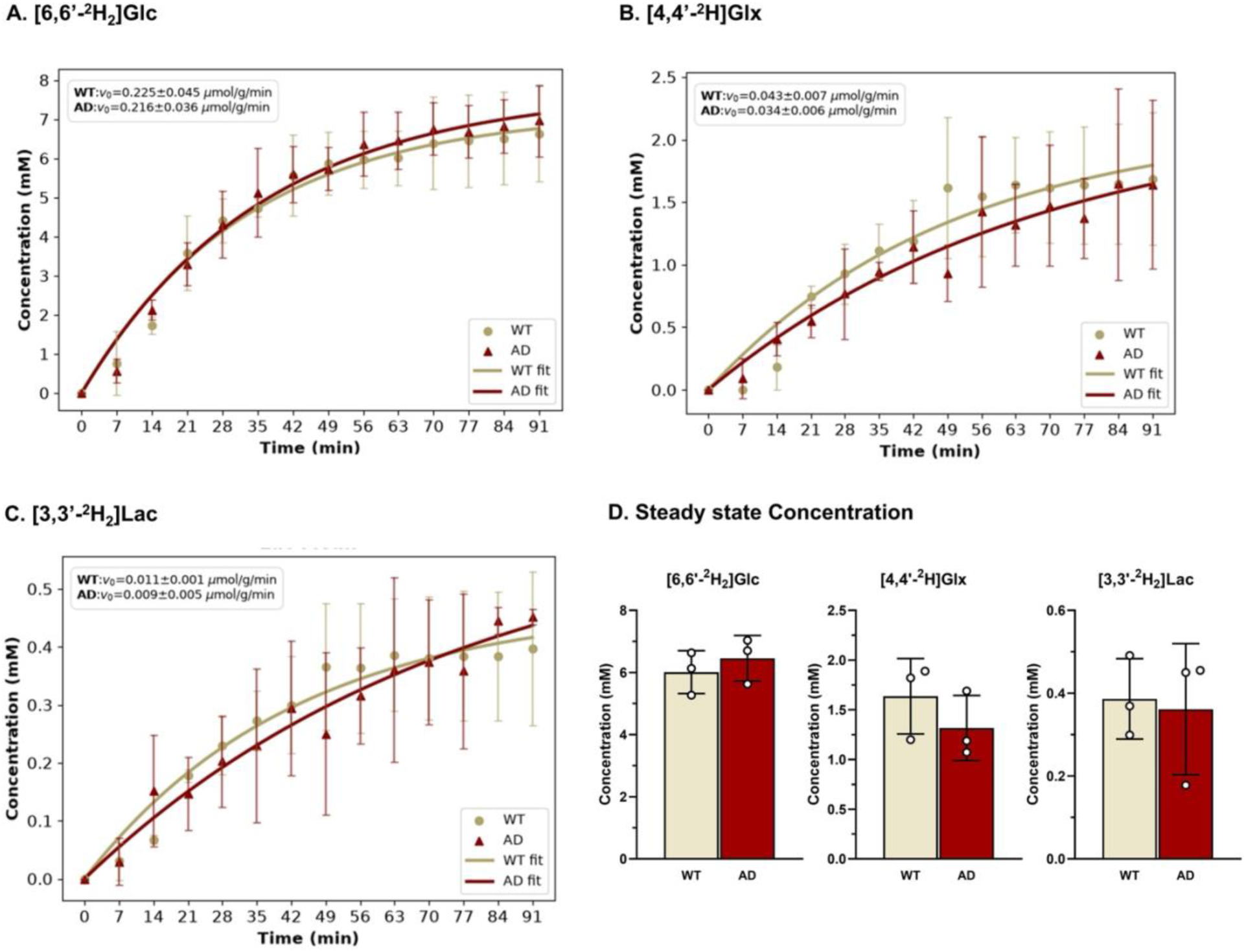
Turnover curves showing concentrations of A. [6,6’-^2^H_2_]Glc, B. [4,4’-^2^H]Glx and C.[3,3’-^2^H_2_]Lac in the brains of wild-type (WT) (ivory circles) and APP ^NL-F/NL-F^ Alzheimer’s disease (AD) mice (red triangles) as a function of time. The symbols represent mean values at each timepoint, and solid lines represent exponential fits to the data. D. Steady state (∼60min) concentrations of [6,6’-^2^H_2_]Glc, [4,4’-^2^H]Glx, and [3,3’-^2^H_2_]Lac represented as means±SD. Abbreviations used are: [2,2’,2’’-^2^H_3_]Ace, [2,2’,2’’-^2^H_3_]acetate; [6,6’-^2^H_2_]Glc, [6,6’-^2^H_2_]glucose; [4,4’-^2^H]Glx, [4,4’-^2^H]glutamate/glutamine; [3,3’-^2^H_2_]Lac, [3,3’-^2^H_2_]lactate.

The steady-state concentration of d_2_-Glx derived from d_2_-glucose showed a slight decrease in AD mice (1.31±0.34 mM) compared to WT mice (1.63±0.38 mM); however, this difference remained statistical insignificant (p=0.2) **(Fig. 5D)**. Similarly, the initial rate of d_2_-Glx formation was comparable with 0.033±0.003 μmol/g/min in AD mice and 0.043±0.006 μmol/g/min in WT mice (p=0.2) **(Fig. 5B)**. Moreover, the steady-state concentrations of d_2_-lactate derived from d_2_-glucose were also comparable between the AD (0.36±0.16 mM) and WT (0.38±0.10 mM, p=0.99) groups. Likewise, the initial rates of d_2_-lactate formation in AD mice (0.009±0.004 μmol/g/min) were not different from those observed in WT mice (0.011±0.001 μmol/g/min, p=0.5; Fig. 5C).

## 4. Discussion

The present study used deuterium magnetic resonance spectroscopy (DMRS) to trace glucose and acetate metabolic fluxes in real time to investigate neuronal- and astrocytic activities, in adenovirus-induced neuroinflammation and in the early stages AD (APP^NL-F/NL-F^) mice. Our findings demonstrate that acute adenovirus administration triggers a pronounced metabolic shift, upregulating astrocytic acetate utilization while simultaneously suppressing glucose-driven neuronal tricarboxylic acid (TCA) cycle activity, typical to neuroinflammatory metabolic signature. Consistently, the pre-symptomatic APP^NL-F/NL-F^ model mirrors this early glial phenotype by exhibiting accelerated astrocytic acetate uptake and a trend towards compromised glucose utilization, thereby establishing a shared compensatory metabolic signature that precedes the onset of severe neurodegeneration.

### 4.1. Impaired Astroglial and Neuronal Metabolism in Adenovirus-Induced Astrogliosis

Adenovirus-induced neuroinflammation led to a stark uncoupling of glial and neuronal metabolism in astrogliosis. Since acetate is selectively taken up by astrocytes via monocarboxylate transporters, primarily through MCT-1 (Hosoi et al., 2009; Pierre et al., 2000) and subsequently converted into acetyl-CoA by mitochondrial acetyl-CoA synthetase (AceCS2) (Fujino et al., 2001). The observed elevation in d_2_-Glx production in AG mice directly reflects enhanced astrocytic TCA cycle flux coupled with increased d_3_-acetate uptake and metabolism. This observation is consistent with emerging evidence that reactive astrocytes undergo metabolic reprogramming, including alterations in monocarboxylate transport and utilization, to support the energetic and biosynthetic demands associated with astrocyte activation and remodeling (Nam et al., 2023). Reactive astrocytes undergo substantial morphological and transcriptional reprogramming (Zamanian et al., 2012), which can be accompanied by altered bioenergetic demands (Robb et al., 2020) to support cellular hypertrophy, increased GFAP synthesis, and the production and secretion of inflammatory mediators.

Interestingly, despite comparable glucose uptake, d_2_-Glx levels derived from d_2_-glucose metabolism in the astrogliosis group were significantly reduced. Although glucose-derived Glx comprises mainly neuronal glutamate and predominantly astrocyte-derived glutamine, the larger neuronal glutamate pool constitutes a substantial proportion of the total Glx signal (Lebon et al., 2002; Deelchand et al., 2009). Enhanced astrocytic TCA cycle activity, as indicated by acetate metabolism, suggests that astrocytic d_2_-glucose utilization and the resulting production of d_2_-glutamine are maintained or elevated in astrogliosis. Because the d_2_-Glx signal represents the combined labeling of glutamate, which is primarily neuronal, and glutamine, which is predominantly astrocytic, the reduction in total d_2_-Glx labeling from d_2_-glucose despite unchanged glucose uptake is unlikely to arise from impaired astrocytic metabolism. Rather, it points to a diminished neuronal contribution, consistent with impaired neuronal oxidative metabolism.

Mechanistically, this phenotype maps onto the dual-tracing metabolic framework described by Nam et al., (2023), which demonstrated that hyper-reactive astrocytes actively upregulate acetate metabolism while simultaneously instigating adjacent neuronal glucose hypometabolism (Nam et al., 2023). Reactive astrocytes may impair neuronal oxidative metabolism through altered metabolic support and inflammatory signaling. Indeed, metabolically reprogrammed ischemic astrocytes have been shown to impair neuronal mitochondrial respiration and promote glutamatergic synapse loss (Borbor et al., 2023). Similarly, inflammatory mediators released by activated glial cells can disrupt neuronal mitochondrial membrane potential and ATP production through mitochondrial calcium dysregulation (Casaril et al., 2022). Notably, glucose-derived lactate labeling remained unchanged in astrogliosis, despite the reduction in glucose-associated Glx. This dissociation suggests that glucose-to-lactate (non-oxidative) labeling is relatively preserved, whereas the oxidative pathway labeling Glx may be differentially affected. However, the relative contributions of neuronal and astrocytic glycolysis remain indistinguishable.

### 4.2. Early Astrocytic Compensation in the Pre-Symptomatic AD Mice

The observed significant increase in brain d_3_-acetate influx in 3-month-old APP^NL-F/NL-F^ mice compared with WT controls suggests altered astrocytic metabolism during early AD progression. Aligned with this observation, findings from a previously published study (Nam et al., 2023) reported increased astrocytic MCT-1 expression and acetate uptake, accompanied by reduced neuronal GLUT3 expression and glucose uptake in 7-14-month-old APP-PS1 and 10-11-month-old 5xFAD mice. Importantly, their human AD investigations also demonstrated increased ^11^C-acetate uptake and reduced ^18^F- FDG uptake in AD-associated brain regions. These findings provide evidence for altered astrocyte-neuron metabolic interactions in both experimental models and human AD patients. Interestingly, in the present study, acetate-derived Glx formation and its steady-state concentration showed a slight though statistically insignificant increase in AD mice. This pattern may suggest an early adaptive response in astrocytic metabolism associated with AD-related pathological stress.

On contrary, glucose-derived Glx and lactate demonstrated modestly decreasing trend while d_2_-glucose uptake remained unchanged in these mice. Although these differences were not statistically significant, the opposing trends between acetate- and glucose-derived metabolites may indicate an early paradigm shift in cerebral metabolism. Previous studies have reported alterations in neuronal and astrocytic metabolism at different stages of AD progression. For instance, Andersen et al., (2017) reported reduced glucose-derived TCA cycle activity in 3-month-old APP-PS1 mice, accompanied by increased glucose-derived lactate labeling (Andersen et al., 2017). Interestingly, acetate-derived metabolic activity remained largely unchanged, suggesting that early metabolic alterations in this model were more pronounced in glucose-dependent pathways. 12-month-old APP-PS1 mice have been shown to exhibit increased astroglial acetate oxidation along with decreased neuronal glucose oxidation, reduced glutamatergic and GABAergic metabolic activity, and impaired neurotransmitter cycling in the hippocampus and cerebral cortex (Patel et al., 2018). These findings provide direct evidence of differential metabolic alterations between astrocytes and neurons within the same AD model. Collectively, these studies suggest that AD pathology may promote increased astrocytic substrate utilization and metabolic reprogramming alongside impaired neuronal glucose metabolism and viability. Such changes may reflect an altered metabolic relationship between astrocytes and neurons, potentially contributing to neuronal and synaptic dysfunction during disease progression.

### 4.3. Acute Inflammatory Insult vs. Early AD Pathogenesis

The metabolic similarities between adenovirus-induced astrogliosis and early-stage AD (APP^NL-F/NL-F^) mice suggest that adenovirus induced astrogliosis model recapitulates important aspects of the early metabolic alterations associated with AD.

Both models exhibit increased astrocytic acetate handling and evidence of altered glucose metabolism, with these changes more pronounced in adenovirus-induced astrogliosis. Importantly, the metabolic phenotype observed in the present astrogliosis model extends beyond a general inflammatory response, as it captures the divergence between astrocytic acetate-associated metabolism and neuronal glucose-associated metabolism also suggested by the early APP^NL-F/NL-F^ findings. Therefore, adenovirus-induced astrogliosis may provide a useful complementary model for investigating early AD-associated metabolic disturbances, particularly the mechanisms underlying altered astrocytic substrate utilization and its relationship with neuronal glucose metabolism. Although it does not fully recapitulate the complexity of AD pathology, its ability to recapitulate selected metabolic features of early AD makes it a potentially valuable in identifying and testing therapeutic strategies targeting early metabolic dysfunction in AD.

## 5. Conclusions

Collectively, these findings suggest that adenovirus-induced astrogliosis provides a useful complementary model for investigating early AD-associated metabolic dysfunctions. The more pronounced metabolic phenotype offers an opportunity to explore the mechanisms underlying altered astrocytic substrate utilization and its potential consequences for neuronal metabolism, which remain relatively subtle in pre-symptomatic APP^NL-F/NL-F^ mice. Although this model does not recapitulate amyloid pathology, it may help identify metabolic mechanisms that link neuroinflammation to the early disruption of neuron–astrocyte metabolic interactions in AD.

## Acknowledgements

This project was supported by the National Institute on Aging grants R01AG091760, R01AG063869, and RF1AG087306, and the National Institute of Biomedical Imaging and Bioengineering of the National Institutes of Health through Grant number P41EB029460.

## Notes

### Competing Interest Statement

The authors have declared no competing interest.

